# Evaluation of stress tolerance in IR64 rice near-isogenic lines carrying *SUB1* and *DRO1* genes

**DOI:** 10.1101/2024.09.11.612414

**Authors:** Ibrahim Soe, Nguyen Thi Thu Hang, Emmanuel Odama, Rael Chepkoech, Taiichiro Ookawa, Abdelbagi M Ismail, Jun-Ichi Sakagami

## Abstract

Flooding and drought significantly reduce rice yield in rainfed environments. Rice varieties that tolerate complete inundation for up to two weeks carry the *SUB1A* gene, while those enduring deeper water conditions for weeks or months have the *SK1* and *SK2* genes. Conversely, the *DRO1* gene, responsible for deep-rooting, helps in water acquisition under drought. In this study, we compared the growth of NIL-SUB1DRO1 a novel rice genotype with an IR64 background regarding its dual tolerance to submergence and drought. Additionally, we assessed its recovery capacity after exposure to stress. Sixteen and thirteen-days old seedlings of three genotypes (Experiment 1-1 and 2-1) and fourteen-days old seedlings of the two genotypes (Experiment 1-2 and 2-2) were tested under submergence and drought stress in a greenhouse experimental condition respectively. Seedlings were submerged for 10 and 7 days and then allowed to recover for 10 and 7 days respectively. In the drought experiment, seedlings underwent 29 days of drought (Experiment 2-1) and 18 days of drought, followed by 10 days of rewatering (Experiment 2-2). Growth parameters were measured before and after treatment, 4 days after submergence in experiment 1-1 and after the recovery periods. Submergence and drought adversely affected growth and performance. Shoot elongation in submerged plants was reduced by 29.2% for NIL-SUB1DRO1 compared to IR64. Accelerated shoot elongation of IR64 negatively affected its recovery. Chlorophyll content and maximum fluorescence of IR64 were significantly lower than other genotypes after 10 and 7 days of complete submergence. Ten days after recovery in experiment 1-1 the Chlorophyll content and maximum fluorescence of IR64 were not significantly different compared to other genotypes. Seven days after of recovery in experiment 1-2, NIL-SUB1DRO1 had significantly higher chlorophyll content and maximum fluorescence than IR64. After 29 days of drought the tiller number and leaf area of IR64 was lower than other genotypes (Experiment 2-1), while in Experiment 2-2 during drought treatment and recovery, NIL-SUB1DRO1 had greater relative water content, leaf water potential, leaf area, SPAD value, dry weights of shoots and roots, root length, surface area and volume compared to IR64. stomatal conductance of IR64 was higher than NIL-SUB1DRO1 during drought, leading to greater water loss and reduced growth during recovery. NIL-SUB1DRO1 absorbed and retained water more effectively under dry conditions. NIL-SUB1DRO1 and NIL-SUB1 is tolerant to submergence and NIL-SUB1DRO1 and NIL-DRO1 to drought, with no negative effects from combining these genes in modern rice varieties for rainfed lowlands.

## INTRODUCTION

Rice is a primary food source worldwide, providing food and livelihood security to half of the global human population (Samal *et al*., 2018). By 2035, rice production needs to increase by 26% to feed the growing population (Seck *et al*., 2012). As a semiaquatic crop, rice faces various biotic and abiotic stresses due to different climatic, hydrologic and edaphic conditions. The increasing incidence extreme of abiotic stresses due to climate change is a significant constraint to meeting rising food demand and achieving global food security (Lesk *et al*., 2016). Approximately 30% of the world’s rice (*Oryza sativa*) is grown at low elevations and depends on rainfall (Bailey-Serres *et al*., 2010). Rainfed farming reduces groundwater depletion, water pollution and soil salinisation, which are often associated with controlled irrigation systems. However, rainfed fields are susceptible to flooding and drought due to inadequate water management. These two abiotic stresses are the most prevalent factors reducing rice yield in rainfed environments, affecting approximately 40 million hectares of rice at various crop stages globally and severely impacting plant growth, development and yield (Barnabás *et al*., 2008). High rainfall over short periods can lead to flooding and low or no rainfall can lead to prolonged dry spells, significantly reducing crop yields (Lobell *et al*., 2011). In some cases, both floods and droughts may occur within the same season at different crop growth stages. In the coming years, rainfed shallow lowland areas will likely experience heavy precipitation during early crop growth stages, resulting in floods followed by dry spells causing drought. Variability in rainfall patterns, intensity and frequency due to climate change are major factors contributing to the unpredictable occurrence of drought and flood conditions.

Submergence leads to low oxygen availability in plants, obstructing aerobic respiration and hindering growth processes. In extreme cases, it can even result in plant death. Photosynthesis, crucial for supplying carbohydrates and O_2_ for aerobic respiration, relies on CO_2_ and irradiation (Setter *et al.,* 1989). However, both CO availability and irradiation are severely reduced by submergence. In water, CO_2_ is limited due to its lower absolute concentration compared toO_2._ For instance, air-saturated water contains 0.268 mol m^−3^ O_2_ but only 0.011 mol m^−3^ CO_2_ at 25°C and it diffuses slowly through the boundary layer (Jackson and Ram, 2003). Additionally, reduced irradiance underwater, especially in turbid floodwater, further inhibits photosynthesis (Setter *et al*., 1987; Das *et al.,* 2009). Certain rice cultivars exhibit distinct growth control strategies to survive submergence. One such strategy is the quiescence syndrome (Colmer and Voesenek, 2009), where shoot elongation is suppressed early in growth to conserve carbohydrates for an extended period (10-14 days) during flash floods This response is regulated by the *SUB1A* gene, a major quantitative trait locus (QTL) responsible for submergence tolerance. FR13A, a highly tolerant rice (*Oryza sativa* L.) genotype from Odisha, India, has been extensively studied for its *SUB1A*-mediated tolerance (Xu *et al*., 2006; Ismail *et al*., 2013; Panda and Sarkar, 2012). Submergence tolerant cultivars can resume growth during desubmergence by utilising stored carbohydrates. Another survival strategy is the escape syndrome, observed in deepwater rice cultivars (Bailey-Serres and Voesenek, 2008; Colmer and Voesenek, 2009)]. These cultivars rapidly elongate leaf sheaths and internodes when faced with prolonged deep flooding. By doing so, they rise above the water surface, aided by the genes *SK1* and *SK2* (Hattori *et al*., 2011).

Drought stress induces a reduction in plant height, leaf area and biomass (Mishra and Panda, 2017). Leaf growth decreases due to limited water potential (Zhu *et al*., 2020), as disrupted water flow from the xylem to adjacent cells decreases turgor pressure, leading to impaired cell development and reduced leaf area in crops (Hussain *et al*., 2018). Characteristics such as leaf rolling and early senescence initiation are notable under drought stress (Anjum *et al*., 2011). Water scarcity adversely affects essential physiological traits of rice, including net photosynthetic rate, transpiration rate, stomatal conductance, water use efficiency, internal CO_2_ concentration, photosystem II (PSII) activity, relative water content and membrane stability index (Zhu *et al*., 2020). Several factors contribute to the decline in photosynthesis, such as stomatal closure, reduced turgor pressure, decreased leaf gas exchange and reduced CO_2_ assimilation, ultimately damaging the photosynthetic apparatus (Gupta *et al*., 2020). Photosynthetic capacities of leaves and water availability in the root zones are pivotal in controlling growth and yield in susceptible rice genotypes under drought conditions (Zhu *et al*., 2020). Drought tolerance refers to the ability of plants to survive under low tissue water content (Kumar *et al*., 2016). Adaptive mechanisms like maintaining higher leaf water potential, better osmotic adjustment or protective actions such as leaf rolling and stomatal closure are associated with plants’ drought tolerance (Tuberosa, 2012). Root characteristics play a crucial role in plant adaptation to drought stress. Crop performance under water stress depends on the root system characteristics of the variety used. Predicting rice production under water stress can be facilitated by considering root biomass (dry) and length (Comas e*t al.*, 2013). Root growth characteristics exhibit diverse responses under water stress, with rice varieties possessing deep and prolific root systems showing better adaptability, especially in deeper soils (Kim *et al*., 2020). Uga *et al*. (2011) found that a rice variety carrying the *DRO1* gene develops deeper and better root distribution under relatively dry upland conditions. They established a significant positive relationship between the *DRO1* gene and panicle weight, suggesting that *DRO1* enhances drought avoidance under natural field conditions with occasional water stress by promoting root elongation to access water and sustain essential physiological processes.

As sessile organisms, plants must cope with submergence and drought stress at some point in their life cycle. In this study, we evaluated the effectiveness of single genes and combined genes to enhance the tolerance to flooding and drought using new rice genotypes called NIL-SUB1, NILDRO1 and NIL-SUB1DRO1, developed in the background of IR64. These genotypes exhibit the ability to recover after stress, thus serving as a resilience mechanism. We further assessed several morpho-physiological characteristics to understand and explain NIL-SUB1, NIL-DRO1 and NIL-SUB1DRO1 response to submergence and drought stress. The parental varieties IR64, FR13A and Kinandang Patong were used to develop these genotypes. The results of this research not only contribute significantly to breeding more resilient rice varieties but also expedite the deployment and dissemination of these varieties to farmers in the future to sustain their productivity under the current worsening climate conditions.

## MATERIALS AND METHODS

### Experiment 1-1, 1-2: Submergence experiment

#### Breeding plant materials and test varieties

IR64-SUB1 (NIL-SUB1) resulted from a cross between IR64 (bred at IRRI, Philippines) and FR13A, while IR64-DRO1 (NIL-DRO1) was developed at NARO in Japan through a cross between IR64 and Kinandang Patong. By crossing IR64-SUB1 and IR64-DRO1, we generated IR64-SUB1-DRO1 (NIL-SUB1DRO1) genotypes with IR64 as their genetic background. NIL-SUB1, NIL-DRO1, NIL-SUB1DRO1 and IR64 are the varieties used in this study.

#### Cultivation conditions

As for the submersion experiment, the main experiment (1-2) was conducted following the preliminary experiment (1-1). First, in Experiment 1-1, we confirmed the growth of NIL-SUB1 and NIL-SUB1DRO1 relative to IR64 for the maximum quantum yield (Fv/Fm) and chlorophyll (SPAD value) in leaves, which is a typical indicator of flood tolerance under complete flood conditions. Next, in Experiments 1-2, in order to clarify the details of the anaerobic response of the NIL-SUB1DRO1 to IR64, the shoot elongation was also confirmed.

During the Experiment 1-1 and 1-2, the day and night temperatures were maintained at 27°C, with a light intensity of 350 μmol m^−2^ s^−1^ for 12 h in the controlled room. We studied two factors: environmental conditions, control and submergence treatments and varietal factors (NIL-SUB1DRO1, NIL-SUB1 and IR64 in Experiment1-1, NIL-SUB1DRO1 and IR64 in Experiment 1-2). The experimental design followed a completely randomised approach, with complete submergence for ten days followed 10 days of recovery in Experiment 1-1, while in Experiment 1-2, seven days of complete submergence followed by seven days of recovery,

In both submergence experiments, seeds from each genotype were placed in Petri dishes containing filter paper moistened with distilled water and left to germinate at 30°C in an incubator under dark conditions for 24 h. The pre-germinated rice seeds were then sown in a commercial soil mix (N:P:K = 0.9:2.3:1.1; pH 4.5–5.2) in the greenhouse. Sixteen-to fourteen-days-old seedlings for Experiment 1-1 and Experiment 1-2. were transplanted into hydroponic sponges (30 mm), which were inserted into seedling trays inside experimental glass containers (45 cm × 45 cm × 60 cm) in a controlled room. A hydroponic solution (Kyowa Corporation) was maintained at 4.5 cm from the container base, the same height as the seedling tray and plant stem base, to acclimate the seedlings for four days in both experiments. The hydroponic solution was adjusted to pH 5.5. It contained 80 mg L^−1^ N, 76 mg L^−1^ P, 188 mg L^1^ K and other minor elements (Tada *et al*., 2014).

The complete submergence treatment was applied after the four days of acclimatisation by removing the hydroponic solution and supplying tap water to the transparent container box, reaching 45 cm above the plant shoot. For plants in the control treatment, tap water was maintained up to the stem base inside the transparent box. The submergence treatment lasted for ten days and 10 days of recovery in Experiment 1-1, while in Experiment 1-2, seven days followed by another seven days of recovery period during which the tap water was replaced with hydroponic solution up to the stem base for plants that received submergence and control treatments. Measurements of different variables were conducted before submergence, after submergence, after recovery periods in both experiments and after four days of complete submergence in Experiment 1-1.

#### Measurements of variables

Shoot length was measured from the base of the stem to the highest shoot tip using a ruler. The maximum quantum yield was measured after at least 2 h in the dark by clipping the chlorophyll fluorescence equipment (AquaPen-P AP-P 100, PSI, Czech Republic) onto a newly developed leaf of the sampled rice plant. A chlorophyll metre (SPAD-502, Konica Minolta Corporation, Japan) estimated the chlorophyll content (SPAD value) in leaves, with an average of three measurements taken on the upper part of the newly fully expanded leaf.

### Experiment 2-1, 2-2: Drought experiment Cultivation conditions

Similarly, Experiment 2-2 aimed to evaluate useful traits associated with drought tolerance using the new rice genotypes NIL-SUB1DRO1 compared to IR64 and assess their capacity to recover after stress release following preliminary Experiment 2-1. The Experiment 2-1 evaluated on tiller number and leaf area, the Experiment 2-2 evaluated photosynthetic traits and plant dry matter production.

The research was conducted in a greenhouse with mean temperature of 28℃ and a humidity of 55.2% in Experiment 2-1, while in Experiment 2-2 mean temperature was 25.4℃ and a humidity of 58.1%. Approximately 12 h of daylight was provided during the experimental periods.

The experiments involved two water status: drought treatment and a well-watered control. Seeds from these genotypes were placed in petri dishes containing filter paper moistened with distilled water and left to germinate at 30°C in an incubator under dark conditions for 48 h. The germinated seeds were carefully selected and directly sown into randomised PVC (polyvinyl chloride) pipes with an inner diameter of 8 cm and a height of 40 cm. The pipes were divided into two 20-cm layers: the A-layer (above 0–20 cm) and the B-layer (below 20–40 cm). Three seeds were initially sown per pipe and later thinned to one after five days. Each pipe was filled with a mixture of 2.6 kg of commercial soil (pH 4.5-5.2) and sandy soil in a 1:1 ratio for Experiment 2-1 and 1:4 ratio in Experiment 2-2. Additionally, 5 g of balanced compound fertiliser (N, P and K, 8-8-8) was added to each pipe.

The drought treatment began by stopping irrigation 13 days after transplanting in Experiment 2-1 (when seedlings were at 3.7–4.1 leaf age) and continued for 29 days. In contrast, daily irrigation was maintained for plants in the control treatment. In Experiment 2-2, drought treatment started 14 days after transplanting (when seedlings were at 3.7–4.1 leaf age) and lasted for 18 days. After the drought period in Experiment 2-2, daily irrigation resumed for both treatments, lasting for an additional 10 days as recovery period. Observations of variables were conducted before drought treatment and after drought treatment in both experiments and after the recovery period in Experiment 2-2.

#### Measurement of variable

In the greenhouse, temperature and humidity were monitored using the long-range wireless connection logger telemoni TML2101-A (AS ONE Corporation, Japan). Soil moisture sensors placed at a depth of 10 cm provided continuous measurements of the soil moisture status in each pipe. Data were recorded using a ZL6 Basic Datalogger (Metre Group, Inc., Pullman, WA, USA) at 60-min intervals throughout the experiment. Stomatal conductance (gs) was measured daily on the newly developed leaf using a porometer (AP4, Delta-T Devices, Cambridge, UK) between 9:00 a.m. and 2:00 p.m. Shoot biomass and leaf area were determined by cutting the shoot and separating leaves and stems. Leaf area was quantified using digital image analysis software (LIA32, developed by Kazukiyo Yamamoto, Nagoya University). After measuring leaf area, leaves and stems were oven-dried at 80°C for 48 h. to a constant weight before determining shoot dry weight. Roots from each layer were washed and stored in 70% ethanol at 4°C for two weeks before root scanning. Root samples from each soil layer were scanned at 6400 dpi (EPSON XT-X830, Epson America Inc., Los Alamitos, CA, USA). The scanned images were analysed using an image analysis system (WinRHIZO, Regent Instruments Inc., QC, Canada) with a pixel threshold value ranging between 165 and 175 to assess root length, surface area and volume. After root analysis, root samples were oven-dried at 80°C to a constant weight, following the same process as with the leaves and stems, to determine root dry weight. The number of tillers was obtained by physically counting the culms. Relative water content (RWC) was measured using fully expanded leaves between 9:00 a.m. and 2:00 p.m. Sampled leaves (5 cm length) were immediately weighed to determine leaf fresh weight (FW). They were then hydrated to full turgidity for four hours under normal room and light conditions. After hydration, the samples were blotted dry of surface water with tissue paper and weighed to obtain turgid weight. Finally, the samples were oven-dried at 80°C for 24 h and weighed to determine dry weight (DW). RWC was calculated using the following formula (Barrs and Weatherley, 1962).

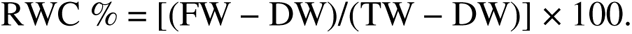

Leaf water potential was measured using a pressure chamber (WP4-T, METRE Group Inc., USA) between 9:00 a.m. and 2:00 p.m. Shoot length and SPAD values were assessed following the same procedure as in Experiment 1-1 and 1-2.

### Data analysis

The data were analysed using a two-way analysis of variance. If significant differences were found, the least significant difference (LSD) test was performed at *p* = 0.05.

## RESULTS

### Effect of submergence on rice plant growth condition

#### Experiment 1-1

Genotype and environment or submergence condition interaction was significant 10 days after complete submergence for SPAD value and Fv/Fm (Table 1). The interaction was also significant for SPAD value 4 days after complete submergence. The SPAD value and Fv/Fm of IR64 decreased significantly after 10 days of submergence compared to NIL-SUB1 and NIL-SUB1DRO1. The SPAD value of IR64 decreased by 47.3% compared NIL-SUB1 and 45.3% for NIL-SUB1DRO1. There was no significant difference between these genotypes before submergence, 4 days after submergence and 10 days after desubmergence. *Experiment 1-2*

**Table 1.** Chlorophyll (SPAD value) and maximum quantum yield (Fv/Fm) of submergence.

| SPAD value |  | 0 DAS | 4 DAS | 10 DAS | 10 DAR |
| --- | --- | --- | --- | --- | --- |
| Control | IR64 | 32.5 ± 2.24 a | 19.8 ± 4.03 a | 11.7 ± 3.53 a | 11.7 ± 4.14 a |
|  | NIL-SUB1 | 30.4 ± 1.87 a | 18.3 ± 1.48 a | 15.2 ± 4.48 a | 12.4 ± 3.40 a |
|  | NIL-SUB1DRO1 | 32.1 ± 1.35 a | 20.3 ± 2.52 a | 11.4 ± 2.89 a | 10.2 ± 3.89 a |
| Submergence | IR64 |  | 19.9 ± 4.61 A | 9.8 ± 3.29 B | 12.2 ± 5.87 B |
|  | NIL-SUB1 |  | 20.6 ± 3.56 A | 18.6 ± 2.88 A | 14.1 ± 5.02 A |
|  | NIL-SUB1DRO1 |  | 20.6 ± 1.80 A | 17.9 ± 3.79 A | 10.7 ± 3.92 A |
| ANOVA (p-value) | T |  | 0.517ns | 0.019* | 0.351ns |
|  | V |  | 0.902ns | <0.001*** | 0.035* |
|  | T*V |  | 0.034* | 0.005** | 0.974ns |
| Maximum quantum yield (FvFm<sup>-1</sup>) |  |  |  |  |  |
| Control | IR64 | 0.772 ± 0.018 a | 0.811 ± 0.023 a | 0.778 ± 0.019 a | 0.796 ± 0.016 a |
|  | NIL-SUB1 | 0.767 ± 0.015 a | 0.810 ± 0.009 a | 0.789 ± 0.003 a | 0.787 ± 0.008 a |
|  | NIL-SUB1DRO1 | 0.768 ± 0.018 a | 0.807 ± 0.008 a | 0.792 ± 0.017 a | 0.788 ± 0.008 a |
| Submergence | IR64 |  | 0.802 ± 0.012 A | 0.711 ± 0.041 B | 0.791 ± 0.013 A |
|  | NIL-SUB1 |  | 0.788 ± 0.011 A | 0.766 ± 0.014 A | 0.788 ± 0.006 A |
|  | NIL-SUB1DRO1 |  | 0.783 ± 0.044 A | 0.767 ± 0.016 A | 0.784 ± 0.007 A |
| ANOVA (p-value) | T |  | 0.003*** | <0.001*** | 0.087ns |
|  | V |  | 0.14ns | <0.001*** | 0.033* |
|  | T*V |  | 0.523ns | 0.015* | 0.186ns |

The interaction of genotype and environment significantly influenced all parameters collected after submergence and recovery, except root DW after submergence and shoot DW after recovery (Table 2). There was no significant difference in shoot length between submerged IR64, control IR64 and NIL-SUB1DRO1 after submergence during the experiment (Figure 2). While the shoot length of IR64 increased significantly (29.2%) when submerged, that of NIL-SUB1DRO1 did not change significantly (8.7%, ns). The slow growth characteristic conferred by *SUB1A* under submergence was evident in the limited change in shoot length of NIL-SUB1DRO1, seemingly independent of the presence of DRO1. After 7 days of complete submergence, chlorophyll content (SPAD value) for IR64 decreased significantly (61.5%), whereas that for NIL-SUB1DRO1 decreased by only 12.7% (not significant) (Figure 3). Although there was no significant difference between control plants and submerged NIL-SUB1DRO1 in SPAD readings after seven days of recovery from submergence, the SPAD values for NIL-SUB1DRO1 were significantly higher than those for submerged IR64. Additionally, chlorophyll fluorescence (Fv/Fm) did not differ significantly between control plants and submerged NIL-SUB1DRO1, while submerged IR64 exhibited significantly lower Fv/Fm (0.63) compared to the control plants (Figure 4).

**Table 2.** Analysis of variance (ANOVA) of submergence Experiment 1-2.

| Parameter | Collection date | Variety | Environment | Interaction (G X E) |
| --- | --- | --- | --- | --- |
| Shoot length (cm) | 0 DAS | ns | ns | ns |
|  | 7 DAS | *** | *** | *** |
|  | 7 DAR | *** | *** | *** |
| SPAD value | 0 DAS | ns | ns | ns |
|  | 7 DAS | *** | *** | *** |
|  | 7 DAR | *** | *** | *** |
| Maximum quantum yield (FvFm<sup>-1</sup>) | 0 DAS | ns | ns | ns |
|  | 7 DAS | *** | *** | *** |
|  | 7 DAR | *** | *** | *** |

\* Significant at $p < 0.05$ , \*\* significant at $p < 0.01$ , \*\*\* significant at $p < 0.001$ ns = not significant, O DAS = 0 days after submergence, 7 DAS = 7 days after submergence and 7 DAR = 7 days after recovery.

### Effect of drought on rice growth condition

#### Experiment 2-1

The soil water content at the start of the drying treatment, when irrigation was stopped, averaged 0.189 (m^3^/m^3^). On the other hand, the soil water content on the last day of the drought treatment, 29 days after the stop of irrigation, decreased to an average of 0.069 (m^3^/m^3^). Twenty-nine days of drought significantly reduced the tiller number and leaf area of IR64, NIL-DRO1 and NIL-SUB1DRO1 (Figure 5 and 6). The leaf area of IR64, NIL-DRO1 and NIL-SUB1DRO1 decreased by 82.6%, 62.25 and 72.05% respectively, while their tiller number decreased by 42.8%, 25.0% and 29.4% respectively compared to control plants. IR64 was significantly lower than NIL-DRO1 and NIL-SUB1DRO1 and no significant between NIL-DRO1 and NIL-SUB1DRO1 for tiller number under drought condition. The leaf area of drought IR64 was lower than drought NIL-SUB1DRO1 and significantly lower drought NIL-DRO1 after 29 days of drought condition.

#### Experiment 2-2

The soil moisture status of IR64 and NIL-SUB1DRO1 is presented in Figure 1. Under control conditions, there was no change in soil moisture content for both genotypes. However, on the second day after ceasing irrigation (drought treatment), the soil moisture content of both IR64 and NIL-SUB1DRO1 began to decrease. By the end of the 18-day drought treatment, it had decreased by 53.9% for IR64 and 53.5% for NIL-SUB1DRO1. During the recovery period, soil moisture levels increased similarly for both genotypes.

The morpho-physiological performance of both genotypes was significantly influenced by the interaction of genotype and environment (Table 3). Notably, there was a significant difference in shoot length between IR64 and NIL-SUB1DRO1 after the drought and recovery periods (Figure 7). While IR64’s shoot length increased by 19.6%, NIL-SUB1DRO1 exhibited a more substantial increase of 44.3% during the drought period. Drought stress led to decreased chlorophyll content (SPAD value) in both NIL-SUB1DRO1 and IR64, with reductions of 8.1% and 26.2%, respectively, after 18 days of drought compared to the control. Interestingly, NIL-SUB1DRO1 maintained significantly higher chlorophyll content after recovery from drought than IR64. Stomata play a crucial role in drought tolerance by controlling CO_2_ uptake and transpiration rates. To explore stomatal responses under drought stress and during the recovery period, stomatal conductance (gs) was measured (Figure 9). Although NIL-SUB1DRO1 exhibited slightly lower stomatal conductance than IR64 during the drought period, this difference was not significant. Furthermore, there was no significant difference in stomatal conductance between the two genotypes during the recovery stage or under control conditions, suggesting that neither of the two genes significantly influenced stomatal responses.

**Table 3.** Analysis of variance (ANOVA) for the drought Experiment 2-2.

| Parameter | Collection date | Variety | Environment | Interaction (G X E) |
| --- | --- | --- | --- | --- |
| Shoot length (cm) | 0 DAD | ns | ns | ns |
|  | 18 DAD | *** | *** | *** |
|  | 10 DAR | *** | *** | *** |
| SPAD value | 0 DAD | ns | ns | ns |
|  | 18 DAD | *** | *** | *** |
|  | 10 DAR | *** | *** | ** |
| Stomata conductance<br>(mmolm<sup>-2</sup>s<sup>-1</sup>) | 0 DAD | ns | ns | ns |
|  | 18 DAD | *** | *** | ** |
|  | 10 DAR | ns | ns | ns |
| Leaf area (cm<sup>2</sup>) | 0 DAD | ns | ns | ns |
|  | 18 DAD | *** | *** | *** |
|  | 10 DAR | *** | *** | *** |
| Leaf relative water content<br>(%) | 0 DAD | ns | ns | ns |
|  | 18 DAD | *** | *** | *** |
|  | 10 DAR | *** | ** | * |
| Leaf water potential (MPa) | 0 DAD | ns | ns | ns |
|  | 18 DAD | *** | *** | *** |
|  | 10 DAR | *** | *** | *** |
| Shoot dry weight (mgplant<sup>-1</sup>) | 0 DAD | ns | ns | ns |
|  | 18 DAD | *** | ** | ** |
|  | 10 DAR | *** | *** | *** |
| Root dry weight (mgplant<sup>-1</sup>) | 0 DAD | ns | ns | ns |
|  | 18 DAD | *** | * | * |
|  | 10 DAR | *** | ** | ns |
| Root length (cm) | 18 DAD | *** | *** | *** |
|  | 10 DAR | *** | *** | *** |
| Root surface area (cm<sup>2</sup>) | 18 DAD | *** | *** | ** |
|  | 10 DAR | *** | *** | ** |
| Root volume (cm<sup>3</sup>) | 18 DAD | *** | *** | *** |
|  | 10 DAR | *** | *** | *** |

\* Significant at $p < 0.05$ , \*\* significant at $p < 0.01$ , \*\*\* significant at $p < 0.001$ ns = not significant, O DAS = 0 days after drought, 18 DAS = 18 days after drought and 10 DAR = 10 days after recovery.

Drought stress also impacted the leaf area (Figure 10) in Experiment 2-2. IR64 had a significantly lower leaf area compared to NIL-SUB1DRO1 after both the drought and recovery periods. The mean leaf relative water content and leaf water potential of IR64 were significantly reduced under drought conditions compared to the control. Specifically, IR64’s relative water content decreased by 9.3% during the drought, while NIL-SUB1Dro1 only experienced a 2.7% reduction (which was not significant) (Figure 11). Interestingly, IR64 did not recover well after 10 days of recovery in terms of leaf relative water content (Figure 11). Additionally, IR64 exhibited significantly lower leaf water potential after drought (−3.88 MPa) and recovery (−2.22 MPa) compared to NIL-SUB1DRO1 (−2.34 MPa after drought and −1.83 MPa after recovery). Finally, drought stress significantly reduced shoot DW for both NIL-SUB1DRO1 and IR64 (by 41.9% and 53.2%, respectively) and root DW for IR64 (by 65.6%) and NIL-SUB1DRO1 (by 40.0%) compared to control plants (Figure 13a and b).

The ratio (B-layer/A + B-layers) of root morphological characteristics, including total root length, root surface area and root volume, after drought and recovery is presented in Table 4. Significantly higher ratios (B-layer/A + B-layers) were observed for NIL-SUB1DRO1 compared to IR64 after both the drought period and recovery from the drought. However, no significant differences were found between the two genotypes under control conditions for these root morphological characteristics.

**Table 4.** Ratios of root length, surface area and volume after drought and recovery (B-layer/A + B-layers) of Experiment 2-2.

|  |  | Root length (cm) |  | Root surface area (cm<sup>2</sup>) |  | Root volume (cm<sup>3</sup>) |  |
| --- | --- | --- | --- | --- | --- | --- | --- |
|  |  | Control | Drought | Control | Drought | Control | Control |
| 18<br>DAD | IR64 | 0.46 | 0.21 | 0.57 | 0.32 | 0.46 | 0.22 |
|  | NIL-SUB1DRO1 | 0.46 | 0.36 | 0.58 | 0.46 | 0.45 | 0.38 |
|  | T-test | ns | *** | ns | *** | ns | *** |
| 10<br>DAR | IR64 | 0.68 | 0.35 | 0.41 | 0.26 | 0.52 | 0.37 |
|  | NIL-SUB1DRO1 | 0.67 | 0.44 | 0.41 | 0.33 | 0.51 | 0.45 |
|  | T-test | ns | *** | ns | *** | ns | *** |

\*\*\* Significant at $p < 0.001$ , ns = not significant, 18 DAD = 18 Days After Drought and 10 DAR = 10 Days After Recovery.

## DISCUSSION

The significant effect of submergence and genotype interaction was the increase in shoot length observed in IR64 compared to NIL-SUB1DRO1 under submerged conditions (Figure 2) in Experiment 1-2. According to the mechanism of submergence defined by Nagai *et al*. (2010) and Hattori *et al*. (2011), IR64’s elongation represents transient submergence intolerance, while NIL-SUB1DRO1’s relative quiescence reflects tolerance mediated by *SUB1A*. Shoot elongation during submergence, caused by flash flooding, can have adverse effects on survival due to wasted carbohydrates and lodging after desubmergence (Ram *et al*., 2002). NIL-SUB1DRO1, carrying the *SUB1A* gene, limits plant elongation during submergence, allowing for greater shoot dry matter accumulation compared to IR64 (Nurrahma *et al*., 2021a). In Experiment 1-1 and 1-2, we assessed the efficiency of PSII in both IR64, NIL-SUB1 and NIL-SUB1DRO1 by measuring their Fv/Fm after submergence and recovery. Chlorophyll fluorescence (Fv/Fm) serves as an effective indicator of rice submergence tolerance (Sone *et al*., 2012). The significant decline in chlorophyll fluorescence observed in submerged IR64 after 10 and 7 days under water likely reflects reduced PSII activity, disorganisation of the photosynthetic apparatus and decreased light intensity and oxygen levels in floodwater (Panda and Sarkar, 2012). Photoinhibition damage occurs in response to environmental stress, leading to decreased solar energy conservation during photosynthesis. While both sensitive and tolerant genotypes experience reductions in chlorophyll concentrations during submergence, the effects are relatively greater in sensitive genotypes (Ella *et al*., 2003; Singh *et al*., 2014). Leaf chlorosis, observed in the sensitive variety, IR64 under both submergence experiments, further reduces photosynthesis. Additionally, submerged rice may experience reduced photosynthetic rates due to declining light intensity and gas diffusion rates (Nurrahma *et al*., 2021b). Dessougi *et al*. (2022) emphasised that chlorophyll plays a crucial role in light absorption, transformation and energy transmission in plants. Therefore, higher chlorophyll content in submerged plants enhances carbohydrate production and survival chances. Evaluating chlorophyll content can serve as an indicator of potential dry matter production and plant survival (Singh *et al*., 2014; Liem *et al*., 2019).

The recent development of high-density linkage maps has provided tools for dissecting the genetic basis of complex traits like drought tolerance into individual components. These efforts have led to the identification of quantitative trait loci (QTLs) related to drought tolerance components such as osmotic adjustment (Robin *et al*., 2003), stomatal regulation (Price *et al*., 1997), leaf water status and root morphology (Kamoshita *et al*., 2002). The ability of a plant to recover after drought stress is also crucial. Some researchers suggest that drought recovery ability is more important than drought tolerance (Sarkar and De Datta, 1975). In this study, genotype and environment interactions significantly affected the performance of IR64, NIL-DRO1 and NIL-SUB1DRO1 after 29 days of drought and NIL-DRO1 and NIL-SUB1DRO1 after 18 days of drought and 10 days of recovery, except for root DW and stomatal conductance after recovery. During the drought and recovery periods, IR64’s leaf relative water content, leaf water potential, leaf area and SPAD value were significantly lower than in control plants. According to Fofana *et al*. (2018), the significantly higher leaf area of NIL-SUB1DRO1 and NIL-DRO1 in this study after drought stress could be supported by the leaf’s morphology (flatter leaves and fewer dead leaves), suggesting a higher assimilatory surface for light capture and transpiration. The higher stomatal conductance observed in IR64 during the drought period could have predisposed the variety to a change in water balance through increased water loss under reduced supply. Under water-limited environments, a plant’s initial response is to prevent a decline in water content by balancing water uptake and water loss rates, a stress avoidance strategy (Verslues *et al*., 2006). Stomatal closure is the immediate and short-term mechanism plants employ in response to potential water loss (Oliver *et al*., 2010). Stomatal conductance is a component of total diffusion that involves mesophyll diffusion. Serraj *et al*. (2008) suggested that rice is better described as a dehydration avoider; thus, the higher stomatal conductance observed for IR64 could be a disadvantage for drought tolerance. Soil moisture stress affects plants’ water status through water potential components (Chakraborty *et al*., 2008). Farooq *et al*. (2012) observed that leaf hydraulic conductance affects responses to drought stress. Reduced cellular turgidity implied by reduced water potential disrupts the structural integrity of leaf cells, causing electrolyte leakage (Farooq *et al*., 2012). Demirevska *et al*. (2008) noted a significant reduction in leaf relative water content with increasing drought severity. Disruption in water balance and changes in leaf morphology negatively affect other physiological processes, such as transpiration and photosynthesis (Fofana *et al*., 2018). This partly explains the significantly lower shoot DW of IR64 plants under soil moisture stress (Figure 13a). *DRO1* likely regulates stomatal function under dry conditions to prevent moisture loss, in addition to enhancing water uptake.

Rice is a shallow-rooted crop susceptible to drought. Root growth becomes restricted during exposure to abiotic stresses, including drought (Kato and Okami, 2010). To adapt to water-limited conditions, rice roots undergo morphological and anatomical changes (Kato and Katsura, 2014). The ability to develop the root system in response to drought stress reflects phenotypic plasticity (Dien *et al*., 2017). According to Yoshida and Hasegawa (1982), since plants acquire water from the soil, root growth is crucial for drought stress resistance. O’Toole and Chang (1979) found that rice varieties with longer and thicker roots were more drought-tolerant than those with shorter and thinner roots. In this study, the ratio of deeper to shallower roots (B-layer/A + B-layers) for NIL-SUB1DRO1 was significantly higher than for IR64 after drought and recovery for total root length, root surface area and root volume (Table 4). The study’s results align with previous reports, indicating that larger root systems (deeper roots, higher root surface area and root volume) are important for drought tolerance in rice (Dien *et al*., 2017). The *DRO* gene enhances the development and distribution of deeper roots under relatively dry upland conditions to acquire water and other mineral elements essential for plant growth (Uga *et al*., 2011). Apparently, introgressing both *SUB1* and *DRO1* through breeding will likely enhance rice adaptation in areas affected by both flooding and drought, either within the same season or during different years, as is common in most rainfed lowlands. Combining the two genes does not seem to have negative interactions and is likely expressed independently. Further studies are needed to verify these results in natural farmers’ fields.

## CONCLUSIONS

Submergence and drought are serious abiotic stresses affecting rice survival and growth, significantly impacting high-yielding varieties like IR64. NIL-SUB1DRO1 combines the submergence tolerance gene and the deep-rooting gene in the genetic background of IR64, while NIL-SUB1 and NIL-DRO1 contains submergence tolerance and deep-rooting genes respectively in the genetic background of IR64 as well. NIL-SUB1DRO1 exhibited characteristics associated with submergence stress tolerance, reflected in the quiescence strategy that suppresses the elongation of the above-ground part during flooding and maintains a higher Fv/Fm. NIL-SUB1 also maintained a higher Fv/Fm value during submergence. When subjected to dry soil conditions, NIL-SUB1DRO1 displayed favourable deep-rooting properties, including improved root length, root surface area and root volume in the lower soil layers, NIL-DRO1 also maintained stable tiller number and leaf area. Our data also suggest that under drought conditions, NIL-SUB1DRO1can reduce leaf stomatal conductance to prevent moisture loss. Combining the two genes does not have any apparent negative impacts under either stress condition. The significant interaction between genotype and environment in both experiments demonstrates that these genotypes have a genetic effect in different environments. Future research should confirm the effect on tolerance and resilience conferred by these two genes during early growth stages, adaptation and ultimately grain yield under natural field conditions.

## Author contribution statement

Jun-Ichi Sakagami designed the study; Ibrahim Soe and Nguyen Thi Thu Hang conducted the study; Taiichiro Ookawa provided rice genotypes NIL-SUB1, NIL-DRO1 and NIL-SUB1DRO1; Emmanuel Odama and Rael Chepkoech assisted in data collection and analysis. Ibrahim Soe and Jun-Ichi Sakagami wrote the original draft. All authors participated in review and editing of the manuscript.

## Acknowledgement

We thank Dr. Yusaku Uga of the National Agriculture and Food Research Organization, Japan for providing IR64DRO1 used to breed NIL-SUB1DRO1.

## Conflict of interest statement

The authors have declared no conflict of interest

## Funding statement

This study is carried out with funds provided by the United graduate school of agricultural sciences, Kagoshima university, Kagoshima, Japan.

## Data availability statement

The article includes all data collected during this study. All the data in this study are available upon request.

## Notes

### Competing Interest Statement

The authors have declared no competing interest.

